# Novel neutrophil subsets associated with sepsis, vascular dysfunction and metabolic alterations identified using systems immunology

**DOI:** 10.1101/2022.02.01.478658

**Authors:** U. Parthasarathy, Y. Kuang, G. Thakur, J.D. Hogan, T.P. Wyche, J.E. Norton, J.R. Killough, T.R. Sana, C. Beakes, B. Shyong, N.R. Zhang, D.A. Gutierrez, M. Filbin, D.C. Christiani, A. G. Therien, C.H. Woelk, C.H. White, R. Martinelli

**Author notes:** these authors contributed equally (U. Parthasarathy, Y. Kuang, G. Thakur, J.D. Hogan). current address Flagship Pioneering (J.R. Killough). current address Third Rock Ventures, Boston MA, USA.

## Abstract

Sepsis is a life-threatening condition caused by a dysregulated host response to infection. Despite continued efforts to understand the pathophysiology of sepsis, no effective therapies are currently available. While singular components of the aberrant immune response have been investigated, comprehensive studies linking different data layers are lacking. Using an integrated systems immunology approach, we evaluated neutrophil phenotypes and concomitant changes in cytokines and metabolites in sepsis patients. Our findings identify novel differentially expressed immature neutrophil subsets in sepsis patients. These and other subsets correlate with various proteins, metabolites, and lipids, including pentraxin-3, angiopoietin-2 and lysophosphatidylcholines, in sepsis patients. These results enabled the construction of a predictive model based on weighted multi-omics linear regression analysis for patient classification. These findings could help inform earlier patient stratification and treatment options, as well as facilitate future mechanistic studies targeting the trifecta of surface marker expression, cytokines, and metabolites.

## Introduction

Sepsis is defined as a dysregulated immune response to infection, mounted by the host <u>1,2</u>. This notion is challenged by the fact that, of more than 100 clinical trials testing the hypothesis that modulating endogenous responses to sepsis will improve survival, no new treatments have been delivered in the last 40 years, with broad spectrum antibiotics and fluid replenishment remaining the standard of care <u>3</u>. Many hypotheses have been proposed to explain the lack of new therapies, including poor translatability of animal models, inappropriate patient selection and a limited understanding of the basic molecular and cellular underpinnings of sepsis <u>4</u>. Nevertheless, over the course of the last decade, multiple studies have advanced our basic knowledge of this disease. Because of these studies, we now know that the disease course is complex and heterogeneous and includes an initial hyper-inflammatory phase that varies over time and is mediated by cells of both the innate and adaptive immune systems, various inflammatory cytokines, proteases, metabolites, and lipids. Ultimately, the combined effects of both pro- and anti-inflammatory responses can result in irreversible tissue damage, endothelial dysfunction, organ failure and death5,6.

Several studies have established the roles of apoptotic depletion of immune cells, monocytes, B-cell populations, and T-cell populations in the pathogenesis of sepsis <u>7-9</u>.

Within the immune compartment, it is recognized that neutrophils are crucial components of the innate immune response during sepsis <u>10</u>, responsible for the release of important regulatory factors, phagocytosis of pathogens, and antimicrobial killing via expression of a range of antimicrobial peptides, proteases, and oxidants, and by formation of neutrophil extracellular traps (NETs) <u>11</u>. However, neutrophils can also exert detrimental effects: localized production of reactive oxygen species (ROS) and excessive NETs can contribute to tissue damage, vascular leak and thrombosis <u>12,13</u>.

Neutrophil functions also become dysregulated in sepsis, with altered migration due to dysfunctional interaction with the endothelium, altered actin cytoskeleton, impaired apoptosis, and altered G-protein-coupled receptor/toll-like receptor signaling <u>14-16</u>.

Furthermore, disease progression in sepsis has been shown to be associated with suppression of neutrophil genes encoding mediators of inflammation and immune modulation <u>17</u>. Neutrophils can express pro- and anti-inflammatory cytokines including IFNγ, TNF and IL-6 in response to host factors and pathogen associated molecular patterns (PAMPs), <u>18-22</u>. These cytokines, and others such as IL-1beta, IL-12 and IL-17, comprise the so-called cytokine storm which develops in the early hyperinflammatory state of the disease <u>23,24</u>. IL-10-producing neutrophils have been reported in mice during sepsis.

Recent studies have started to investigate the potential crosstalk between cytokine and metabolites in infection/inflammation. The metabolic activity of host cells can be hijacked by bacteria and viruses to facilitate their replication, driving alteration of intracellular metabolites and dysregulation of metabolic enzymes that can directly and indirectly regulate immune responses <u>25-27</u>. Indeed, dysregulated metabolites have been reported in sepsis <u>28</u>, with a predominant catabolic state leading to the breakdown of carbohydrates, lipids, and proteins reported <u>29-33</u>.

Most studies have focused on characterizing neutrophil responses, cytokines or metabolites as singular data layers in sepsis, and this information taken by itself can be potentially limiting/misleading <u>34</u>. Relating plasma levels of soluble factors and metabolites with cellular profiles of specific immune cell subsets in sepsis patients could not only strengthen predictive findings, but could also help elucidate the mechanisms of disease progression and enable stratification of patient populations. To this end, we used an integrated systems immunology approach, that included in-depth profiling at the cellular (high-dimensional flow cytometry), and molecular levels (plasma cytokines and metabolomics/lipidomics) to further characterize neutrophil signatures in sepsis patients.

## Results

### Clinical cohort information and systems immunology approach

We utilized an integrated analysis approach to characterize the cellular, molecular, and metabolic immune signatures in sepsis. To this end, we performed flow cytometry, cytokine profiling, and metabolomics/lipidomics analyses on sepsis patients (n = 24) at diagnosis and age/gender-matched healthy donors (n = 19) (**Figure 1A**). Clinical information for patient groups and parameters for identification of sepsis are summarized in **Table S1**.

**Figure 1.**
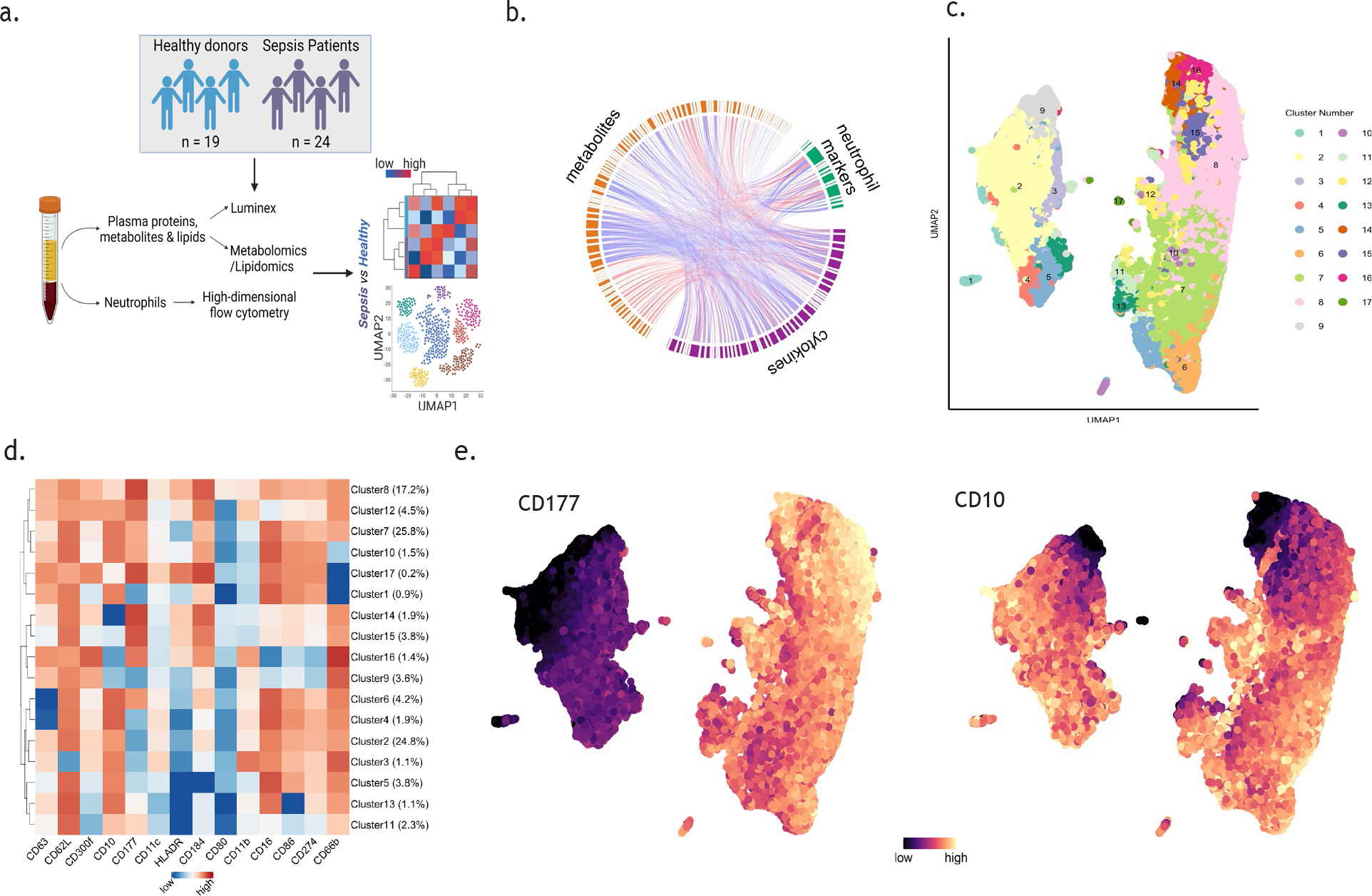
Cell clustering and correlation analysis identifies multiple populations of neutrophils. **(A)** Patient cohort and experimental design used in this study. **(B)** Circos plot showing relationships between features in the three data layers (neutrophil surface marker expression, cytokines, and metabolites/lipids); orange connections are between neutrophil markers and metabolites/lipids, purple connections are between neutrophil markers and cytokines, and gray markers are between metabolites/lipids and cytokines. The width of nodes around the circular layout corresponds to the sum of each feature’s degree distribution. **(C)** UMAP plot of neutrophils based on flow cytometry analysis in total cohort (sepsis patients and healthy controls). Each color represents a different cell cluster (identified by FlowSOM), and each dot represents a single cell. **(D)** Hierarchical clustering and heatmap showing differential expression of individual surface markers across 16 clusters in total subject cohort, colored from low median marker expression (blue) to high median marker expression (red). **(E)** UMAP plot of neutrophils colored according to CD10 and CD177 expression for each cell, colored from low marker expression (dark blue) to high marker expression (bright yellow).

To assess how different data layers relate to each other, we measured correlation between features in each of our three data layers. We employed a circos plot to visualize correlations between neutrophil surface markers, cytokines, and metabolites whose false discovery rate (FDR) was less than 0.05 (**Figure 1B**). A high level of correlation was observed between neutrophil cell surface markers, cytokines, metabolites and lipids (**Supplemental File S2**). This finding reflects the association between neutrophils and soluble cytokine/metabolites in the pathophysiology of sepsis.

### Identification of novel subsets of immature and mature neutrophils

To elucidate neutrophil heterogeneity and identify different subsets of neutrophils, we employed a flow cytometry-specific version of the self-organizing map (SOM) algorithm, FlowSOM <u>35</u> to cluster cells based on expression of fourteen surface markers of interest. This analysis was performed on both sepsis patients and healthy volunteers as a single cohort. FlowSOM identified seventeen neutrophil clusters, across two major distinct populations, which we visualized using a uniform manifold approximation and projection (UMAP) <u>36,37</u> (**Figure 1C**). We then calculated the median surface marker expression across all cells for each cluster and generated a heatmap with hierarchical clustering to characterize neutrophil subsets (**Figure 1D**). This allowed us to detect canonical neutrophil populations (i.e., CD16^+^ CD11b^+^ CD66b^+^ neutrophils), canonical immature neutrophil populations (i.e., CD10^-^), as well as novel neutrophil subsets. For example, the most abundant clusters (2, 7 and 8) had a mature phenotype (CD10^+^) and differed in expression of CD80, CD177, CD184, and HLA-DR. Cluster 17 was characterized by a mature phenotype and high expression of CD274 and CD300f.

Together with cluster 8, it may represent a novel immunosuppressive neutrophil population. We also identified novel CD10^-/low^ immature populations of neutrophils (clusters 9, 14 and 16), that differed in expression levels of CD11b, CD16, CD80, CD86, CD177, CD184, CD274 and HLA-DR.

As CD177 appears to be expressed approximately in half of the cluster that were identified (**Figure 1D**), we postulated that this neutrophil marker could be the driver of the large subgroups identified in the UMAP (**Figure 1C**). We were also interested in assessing the distribution of immature neutrophils across these two subgroups. We thus superimposed expression of CD10 and CD177 onto the UMAPs (**Figure 1E**), confirming that neutrophils can be split into two large subgroups characterized by presence or absence of CD177. Furthermore, there are also three distinct subsets of CD10^-/low^ immature neutrophils, two of those in the CD177^+^ cluster and the remaining being CD177^-^. In summary, these analyses revealed multiple distinct subpopulations of neutrophils, some of which have not been observed previously.

### Neutrophil subsets are differentially abundant in sepsis patients compared with healthy donors

Next, we assessed the relative differences in neutrophil subsets between sepsis patients and healthy controls, using surface marker expression across all neutrophils per sample and cluster abundance measurements. In an unsupervised approach, we calculated the median marker expression across all neutrophils to generate a multidimensional scaling (MDS) plot <u>38</u> (**Figure 2A**). Good separation between the two cohorts suggested a distinction between sepsis patients and healthy controls based on neutrophil markers alone. In a supervised approach, we further used linear regression to test which markers were differentially expressed between sepsis patients and healthy controls. We found that 10 of the 14 markers showed statistically significant differences between the two groups: CD10, CD16 and CD86 are downregulated in sepsis, while HLA-DR, CD11b, CD80, CD184, CD63, and CD66b are upregulated in sepsis (**Figure 2B**). Hierarchical clustering of samples demonstrated that median marker expression clearly separated sepsis from control, which indicated that neutrophil surface markers are differentially regulated in the pathological state (**Figure 2C, Figure S1A-B**).

**Figure 2.**
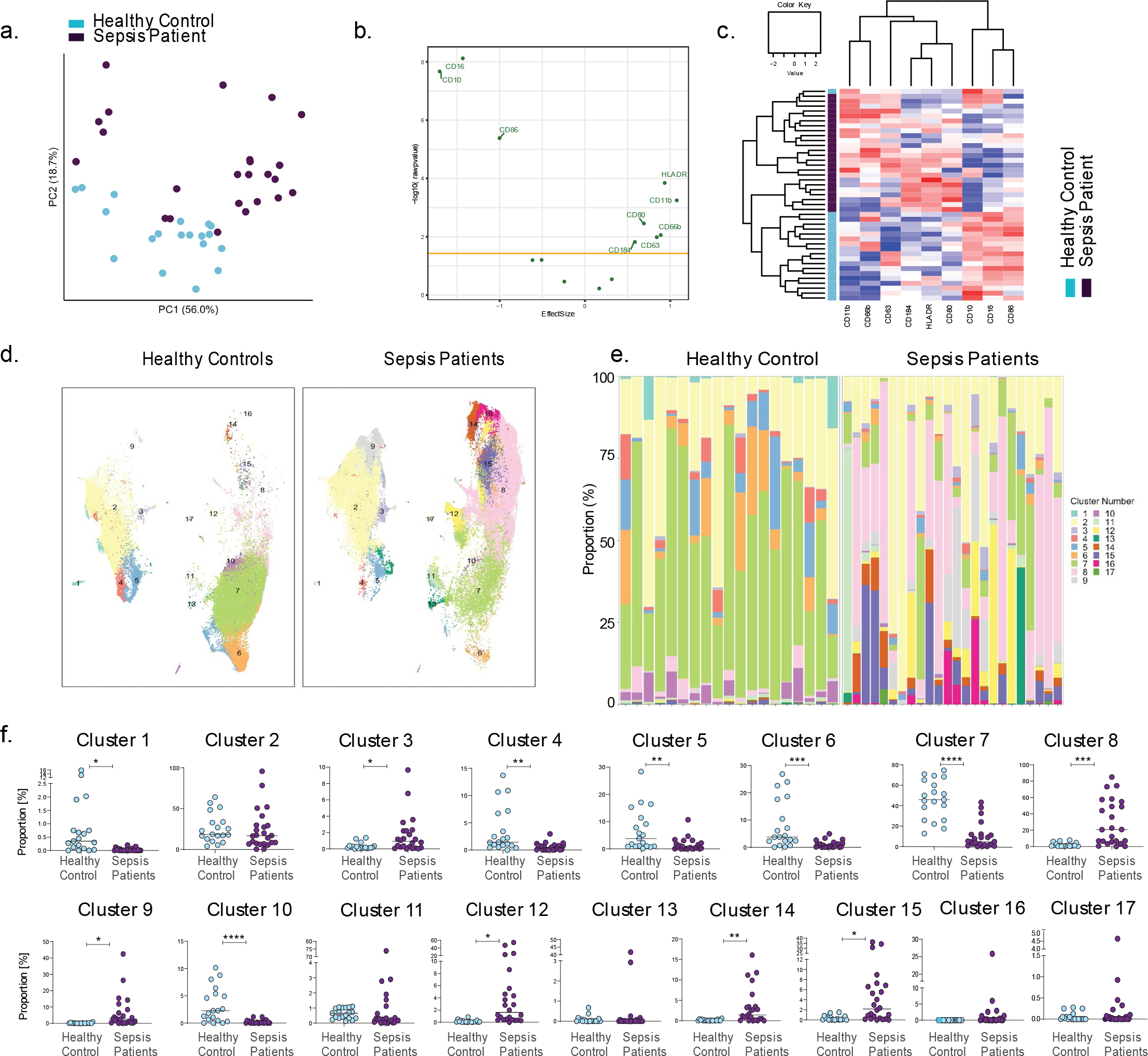
Neutrophil populations display differential marker expression and abundance in sepsis patients compared to healthy controls. **(A)** Multi-dimensional scaling (MDS) plot of median antigen expression in sepsis patients (n=24) and healthy controls (n=22) and showing cohort-specific sample clustering. **(B)** Volcano plot showing effect size and -log10 p-value with the orange line representing the significance cutoff (FDR corrected p-value < 0.05). **(C)** Heatmap of median marker expression across all cells per sample. The top dendrogram represents the similarity between markers while the left dendrogram represents similarity between samples, both of which were determined with hierarchical clustering. The bar on the left indicates sample group with teal (sepsis) and dark purple (control) colors. **(D)** UMAP plot of neutrophil cell clusters in healthy controls and sepsis patients. **(E-F)** Proportion (%) of each cluster in individual sepsis patients/healthy controls **(E)** and between the two clinical cohorts **(F)**. Differential abundance was determined by fitting data to a generalized linear model, and then testing coefficients with a log-ratio for significance (asterisks and FDR adjusted p-value settings) (* = 0.05>FDR>0.01, ** = 0.01 > FDR > 0.001, *** = FDR < 0.001).

While median marker expression suggested differential signals in the two cohorts, UMAPs indicated that specific subpopulations of neutrophils show variations in cluster abundances between sepsis patients and healthy controls (**Figure 2D**). We utilized a generalized linear model to test if clusters were differentially abundant between sepsis patients and healthy controls. We found that clusters 3, 8, and 14 were significantly more abundant in sepsis patients, while clusters 4, 5, 6, 7, and 10 were significantly less abundant in sepsis patients (FDR adjusted p-values ≤ 0.05) (**Figure 2E-F**).

### Novel immature neutrophil subsets are differentially regulated in sepsis

Our results (**Figure 1E**) indicated the presence of three distinct populations of immature neutrophils (CD10^-/low^), largely differentiated by the presence or absence of CD177. As immature neutrophils play a critical role in acute inflammation, we investigated whether they are differentially expressed in sepsis patients and healthy controls. As expected, neutrophils from both healthy donors and sepsis patients clustered accordingly to CD177 expression (**Figure 3A-B**). Within these two clusters, the populations of CD10^-^ immature neutrophils were evident and clearly different in abundance in healthy controls versus sepsis patients. This was confirmed by manual gating on CD10^-^CD16^+^ immature population of neutrophils (**Figure 3Ci**): while in healthy individuals the percentage of immature neutrophils was as expected very low (1.361 ± 1.344 %, mean ± SD), in septic patients it reached levels as high as 70% of all neutrophils (26.46% ± 20.079%, mean ± SD) (**Figure 3Cii**).

**Figure 3.**
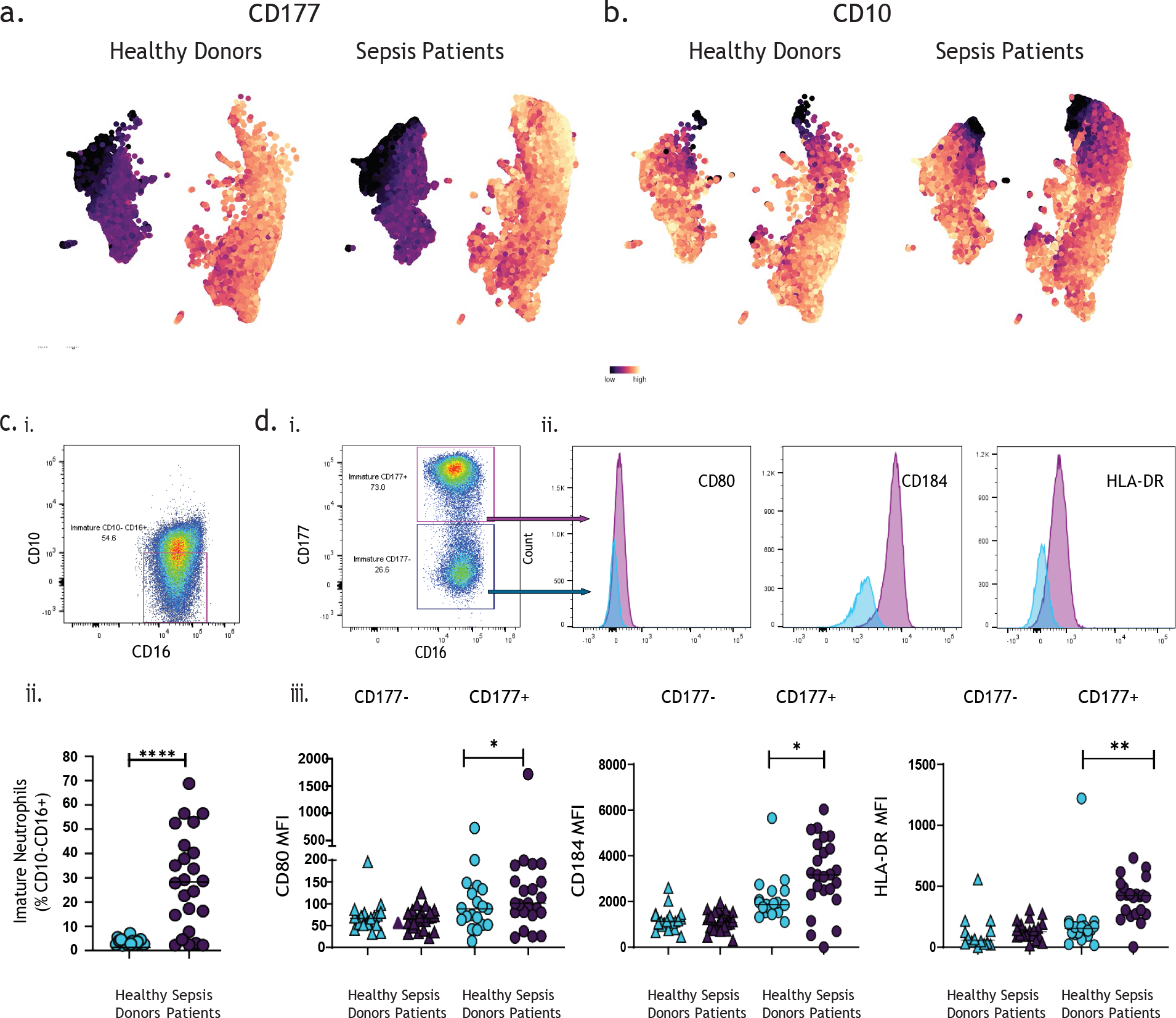
Novel neutrophil subpopulations are differentially expressed in sepsis patients. **(A-B)** UMAP plot of neutrophil cell clusters in healthy controls and sepsis patients colored by CD177 expression **(A)** and CD10 expression **(B)**, colored from low marker expression (dark blue) to high marker expression (bright yellow). **(C)** Selection of immature neutrophils via manual gating on CD10^-^CD16^+^ **(i)** and comparison of this cluster’s percentage in healthy controls vs. sepsis patients. **(D)** Separation of CD10^-^CD16^+^ immature neutrophils by CD177 **(i)**, comparison of CD80, CD184, and HLA-DR expression in the two separated populations **(ii)**, and comparison of CD80, CD184, and HLA-DR median fluorescence intensity (MFI) between healthy controls and sepsis patients in CD177^-^ immature neutrophils and CD177^+^ immature neutrophils **(iii)**. * p<0.5, ** p<0.01, *** p<0.001, t-test.

We also confirmed the presence of the newly identified immature clusters 9 (CD177^-^) and combined 14 and 16 (CD177^+^) (**Figure 1D**) by manual gating, and assessed expression of their key differentiating markers CD80, CD184, and HLA-DR **(Figure 3D, i-iii).** The data show that the two CD10^-^ CD177^+^ immature neutrophils subsets are significantly more abundant (t-test, *p*-value <0.05) in sepsis patients (**Figure 3Diii**). On the other hand, the immature CD177^-^ subset did not display differential expression of CD80, CD184 or HLA-DR, between healthy subjects and septic patients.

### Expression levels of surface markers on neutrophil subsets shows novel correlation with cytokines and other soluble factors in sepsis patients

Next, we evaluated the differential expression of cytokines and vascular-associated factors between sepsis patients and healthy controls. Hierarchical clustering across cytokines and soluble factor levels shows clear separation between the two cohorts (**Figure 4A**), confirming that their dysregulation accompanies sepsis pathology <u>39-41</u>.

**Figure 4.**
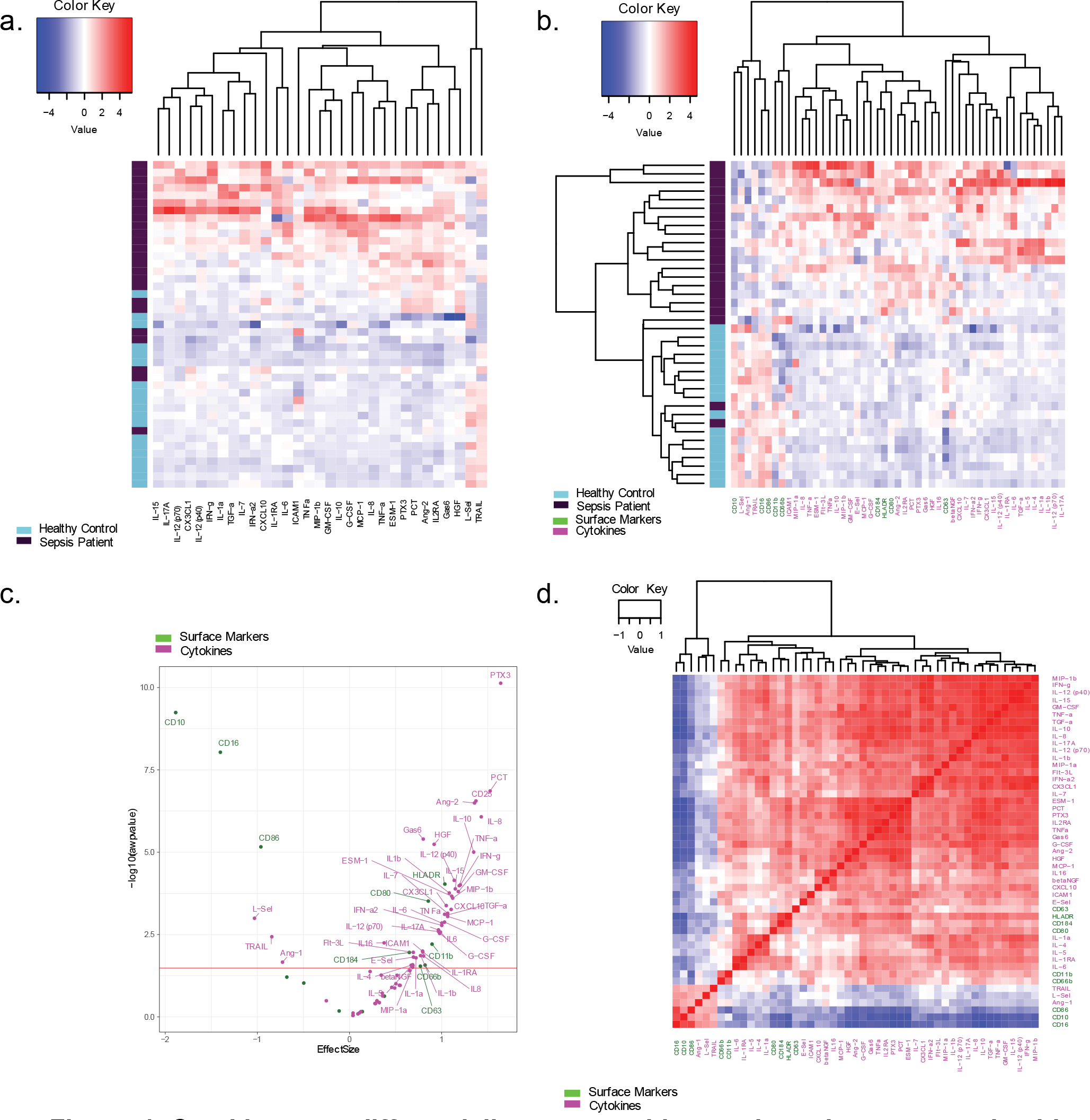
Cytokines are differentially expressed in sepsis patients versus healthy controls and are correlated with specific neutrophils subsets. **(A-B)** Heatmap for significant cytokines only for sepsis patients (n=24) and healthy controls (n=20) **(A)** and heatmap for combined significant cytokines and flow cytometry markers for sepsis patients (n=21) and healthy controls (n=17) **(B)**. Dendrograms on the x and y axes show the unsupervised hierarchical clustering of features and subjects, respectively. The colored bar on the left represents group membership, teal for healthy controls, dark purple for sepsis patients. **(C)** Volcano plot for neutrophil surface markers and cytokines. Processed data from these two data layers were concatenated and a comparison of all surface markers and cytokines isolated from the blood of sepsis patients (n=21) or healthy controls (n=17) is depicted. Markers identified as significant were labeled on the plot (FDR *t*-test < 0.05). The orange horizontal line represents the significance cutoff. Cytokines and surface markers are labelled in magenta and green, respectively. **(D)** Correlogram of significant features obtained from the combination of flow cytometry and cytokine data layers. Pairwise correlation between features is based on Spearman rank correlations. Neutrophil surface markers are depicted in green, and cytokines are depicted in magenta.

An improved separation between healthy controls and septic patients was observed when simultaneously analyzing levels of surface markers measured by flow cytometry and levels of soluble factors measured in plasma (**Figure 4B**).

While upregulation of cytokines is generally observed in the sepsis group, there is notable heterogeneity across individual sepsis patients. For example, although clearly distinguished from healthy donors, not all patients seem to have an increase in TNF-α and IL-6. Also, C-reactive protein, a known marker of sepsis <u>42</u>, was not significantly differentially expressed between healthy subjects and septic patients (data not shown). The highest upregulated factor in sepsis, by effect size (Effect Size = 1.64, adjusted p-value = 5.69e-09) was pentraxin 3 (PTX3); other molecules involved in vascular function, such as endocan (ESM-1), E-Selectin (E-Sel) and Gas6 were also significantly upregulated (FDR p-value ≤ 0.05) in septic patients **(Figure 4C)**. Conversely, decreased levels of L-Selectin (L-Sel) and angiopoietin-1 (Ang-1) were observed in septic patients (**Figure 4C, Figure S3**). To identify relationships between various neutrophil subsets and cytokine expression, we mapped associations both within and between the flow cytometry and cytokine data layers using Spearman’s rank correlation. This analysis showed that CD10 expression in sepsis patients is positively correlated with CD16 and L-Sel, and negatively correlated with PTX3, procalcitonin (PCT), Ang-2, ESM-1, ICAM- 1, and others, indicating a possible role of immature neutrophils in the regulation of endothelial dysfunction and vascular leak (**Figure 4D**).

### Expression levels of surface markers on neutrophil subsets shows novel correlation with metabolites and lipids in sepsis patients

Metabolic alterations have been recognized to be part of the dysregulated response in sepsis <u>43</u>. We therefore extended our analysis to include plasma metabolites and lipids.

To this end, we carried out targeted metabolomics using triple quadrupole (QQQ) liquid chromatography/mass spectrometry (LC/MS). We measured levels of 306 metabolites and lipids in plasma samples from a subset of 12 septic patients and 10 healthy controls for which we had previously generated flow cytometry data. As in the case of cytokines (**Figure 4A**), a heatmap generated by the significantly associated features (t-test) revealed a clear difference between sepsis patients and healthy controls, with the two cohorts almost exclusively clustering with each other (**Figure 5A**). The concentration of several metabolites and lipids, such as lysophosphatidylcholines (lysoPCC16:0, C17:0, C18:0, C18:2), phosphatidylcholines (PC aa C34:4, PC ae C34:2, C34:3, C36:5, C40:1), tryptophan (Trp) and indole propionic acid (IPA), was decreased in sepsis patients compared to healthy controls (**Figure 5A**). On the other hand, the concentration of some metabolites was significantly increased in septic patients compared to healthy controls, for example fatty acids (FA 18:1, 20:1, 18:2, 20:3), putrescine, spermidine, and L-kynurenine, among others (**Figure 5A**). To identify the relationship between various neutrophil subsets and metabolites, we integrated surface markers and metabolite layers, as we did for surface markers and cytokines/soluble factors (**Figure 4B**).

**Figure 5.**
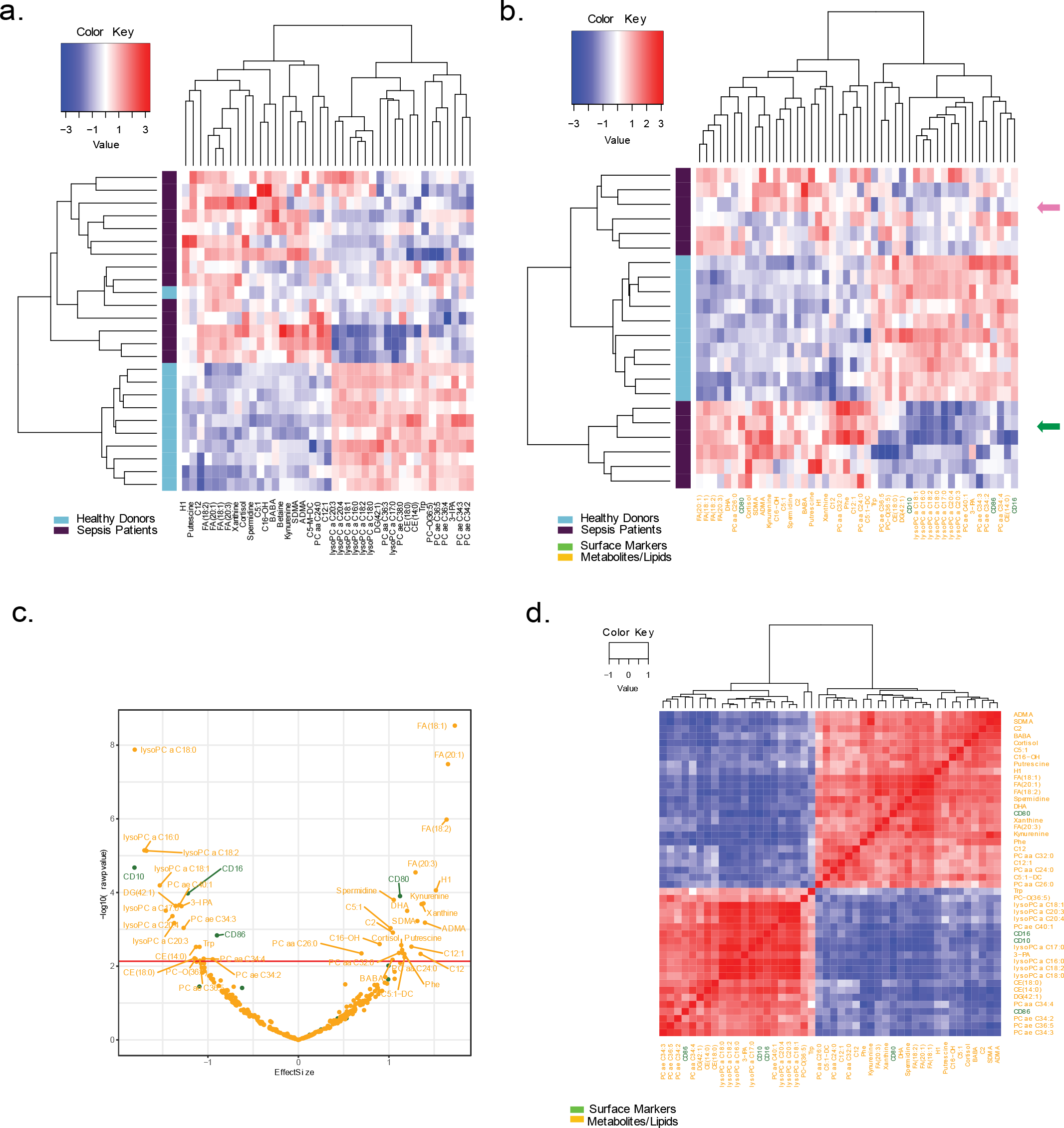
Metabolites and lipids are differentially expressed in sepsis patients versus healthy controls and are correlated with specific neutrophils subsets. **(A-B)** Heatmap for significant metabolites for sepsis patients (n=14) and healthy controls (n=11) **(A)** and heatmap for combined significant metabolite and neutrophil surface markers from sepsis patients (n=12) and healthy controls (n=10) **(B)**. Dendrograms on the x and y axes show the unsupervised hierarchical clustering of features and subjects, respectively. The colored bar on the left represents group membership, teal for healthy controls, dark purple for sepsis patients. **(C)** Volcano plot for neutrophil surface markers and metabolites. Flow cytometry and metabolite data was concatenated for sepsis patients (n=12) and healthy controls (n=10). Markers identified as significant are labeled on the plot (FDR *t*-test < 0.05). The orange horizontal line represents the significance cutoff. All metabolites and surface markers are labeled in orange and green colors, respectively. **(D)** Correlogram of significant features obtained from the combination of flow cytometry and metabolite data layers. Pairwise correlation between features is based on Spearman rank correlations. Neutrophil surface markers are colored green and metabolites are colored yellow.\

Strikingly, the heatmap showed perfect differential clustering of healthy controls from septic patients. In addition, a clear separation between two populations of septic patients was identified (**Figure 5B**). The change in metabolites/lipids was largely homogeneous in the septic group at the bottom part of the heatmap (**Figure 5B**, green arrow). The other septic subgroup, while clearly different from the healthy controls, was heterogenous in terms of metabolite and lipid changes (**Figure 5B**, pink arrow). We next evaluated correlations between surface markers and metabolites/lipids. Several significant (p-value < 0.01) positive and negative (**Figure 5D**) correlations were observed between all surface markers associated with novel immature and mature neutrophil subsets, and metabolite levels. For instance, significant downregulation of CD10 and CD16 and upregulation of CD80 in sepsis neutrophils is correlated with increased production of fatty acids and decreased production of lysoPC, PCaa/PCae species, and 3-IPA (**Figure 5C-D**).

### Predicting novel associations between neutrophil surface markers, cytokines and metabolites in sepsis patients using weighted multi-omics linear regression

Finally, we developed a predictive model for sepsis by examining features in an intersection of subjects in all three data layers: we concatenated the data layers and used the intersection of subjects across all data layers to build a multi-omic classifier with features selected using weighted LASSO regression (see methods for details).

This resulted in an accuracy measure of 0.967±0.039 and an AUROC measure of 0.969±0.040. We then generated a heatmap to visualize the feature expression patterns in relation to the phenotype (sepsis vs. control), and hierarchical clustering demonstrated perfect separation between sepsis and control groups (**Figure 6A**).

**Figure 6.**
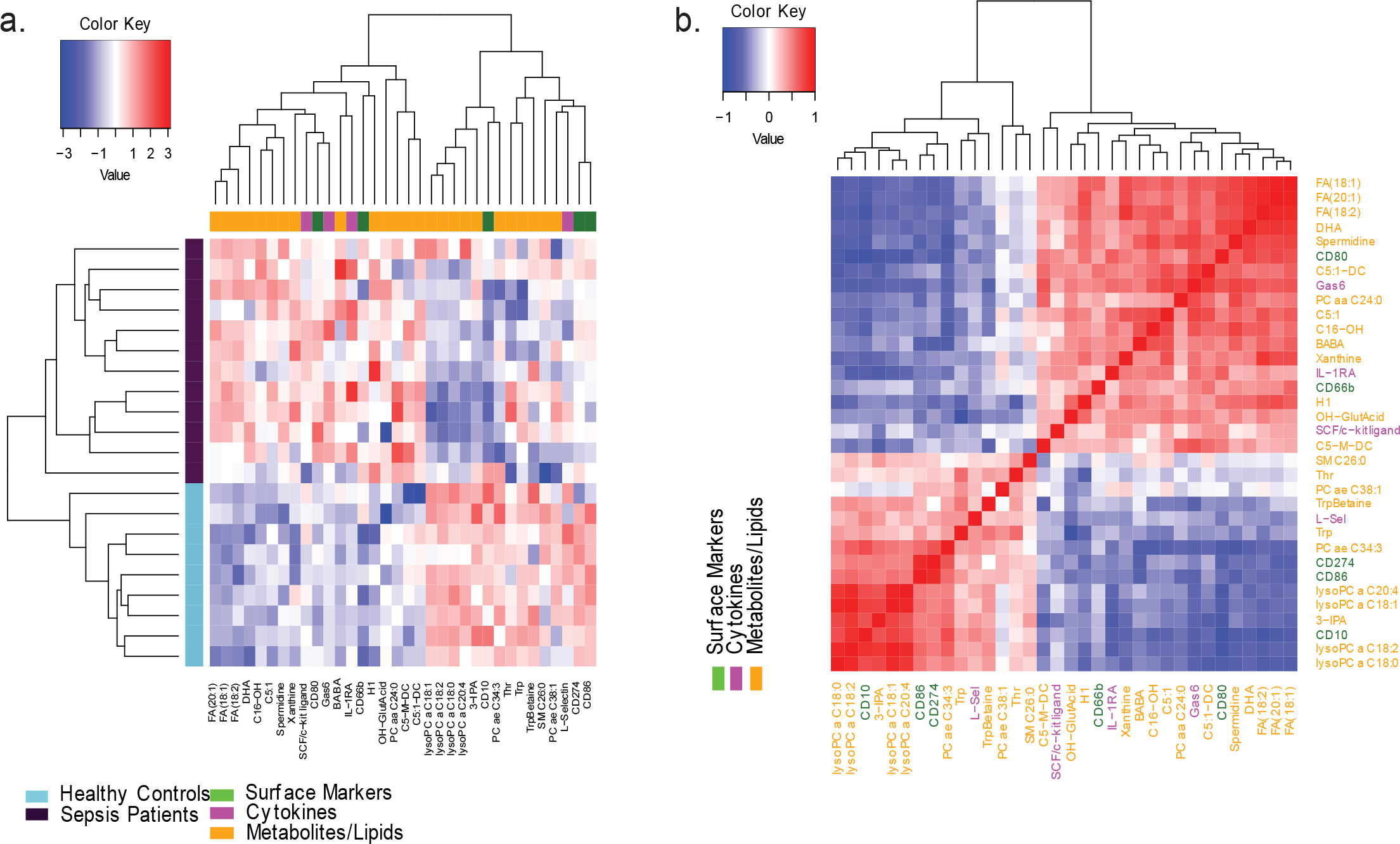
Multivariate approach identifies best classifier between sepsis patients and healthy controls Heatmap for multivariate multi-omics analysis. Processed data from all data layers (flow cytometry, metabolomics, and cytokines) measured from the blood of sepsis patients (n=12) or healthy controls (n=9) are concatenated into a single dataset. Feature selection (employing LASSO regression) was used in nested cross validation framework to obtain the most important set of heterogenous markers to distinguish between sepsis and healthy individuals. Heatmap of expression patterns for all the discriminative features. **(A)** Correlogram showing pairwise Spearman rank correlations between discriminative features **(B)**. Flow markers are colored green, cytokines are colored magenta and metabolites are colored yellow.

CD10 appears again as a critical neutrophil surface marker (**Figures 1-3, Figure 6A-B**), and it is inversely correlated, for example, with fatty acids, as noted previously (**Figure 2C, Figure 6A-B**). Other surface markers that have discriminatory power between sepsis and healthy subjects include CD80, CD86 and CD274. We observed additional features such as Trp, Betaine and OH-Glut Acid that did not have a significant fold-change in previous linear analyses but were important in the non-linear classification of sepsis from healthy subjects (**Figure 6A-B**).

We also noted that each data layer contributed features to the classifier and that there was a higher than expected increase (fisher’s exact test, p = 0.012) in the observed proportion of features selected from the flow cytometry data layer (**Figure S3A**). The proportion of features from the flow cytometry data layer increased from 4.1% to 14.3% in the multi-omics panels and suggested that flow cytometry data was over-represented in the multi-omics classifier and that the flow cytometry data layer contained valuable information to classify sepsis patients.

## Discussion

Despite multiple efforts to characterize the host response and find a clear signature to support development of new therapeutic options, sepsis remains an unmet clinical need. Several groups have characterized immune responses based either on transcriptomics <u>7,44,45</u>, surface protein expression <u>46</u>, or metabolomics <u>33</u>.

Nevertheless, comprehensive approaches that profile the immune response more holistically in sepsis patients are still lacking <u>7,47,48</u>. Specifically, the functional impact of different neutrophil subsets in sepsis is not completely understood <u>46,49</u>. Here, we focused on characterizing neutrophil-driven immune responses and associated pleiotropic effects, by developing an integrative platform including high-dimensional flow cytometry, multiplex bead array for cytokine measurement, metabolomics and lipidomics. Firstly, we confirmed neutrophil heterogeneity **(Figure 1C-D)** and further identified novel subsets of neutrophils. Secondly, we showed how these different subsets are differentially regulated in sepsis compared to healthy controls **(Figure 2-3)**. Lastly, we correlated for the first time the increased presence of immature neutrophils in sepsis patients with endothelial dysfunction and metabolic alterations **(Figure 4-5)**.

High levels of circulating, banded, immature neutrophils (characterized by U-shaped nucleus) is an established, critical feature of the systemic inflammatory response in sepsis <u>50,51</u>. More neutrophils, both mature and immature, are released from the bone marrow to respond to an increase burden of pathogens, and granulopoiesis is enhanced to produce de novo neutrophils. Traditionally, it was thought that immature neutrophils are less competent, with a lower ability to combat infection <u>52</u> and migrate to the site of infection <u>53-55</u>. However, it has recently been shown that the banded immature neutrophil population has a higher phagocytic ability <u>56</u>. This suggests that immature neutrophils may indeed be critical players in managing the infection, but possibly also in exacerbating the immune response.

Rather than relying on nuclear morphology, we defined immature neutrophils by evaluating the levels of the surface marker CD10 <u>49</u>. In this study, we identified two novel subsets of immature neutrophils, CD10^-^CD177^+^ and CD10^-^CD177^-^ (**Figure 1D**) that are differentially regulated in sepsis (**Figures 2C-E**). CD177 function is poorly understood and distinct roles for CD177^+^ and CD177^-^neutrophils subsets have not yet been established. As CD177 is localized to the specific granules for rapid mobilization to the surface upon cell activation, we could speculate that this immature population of neutrophils might drive some of the critical neutrophil functions such as phagocytosis, ROS and NETs production, based on the co-expression of other markers.

We observed significant differential expression of CD184 (CXCR4) and HLA-DR in the CD10^-^CD177^+^ immature neutrophil subset in sepsis versus healthy individuals. CD184 has been known to control the release of neutrophils from the bone marrow and their recirculation as aged senescent neutrophils back to the bone marrow <u>57,58</u>. Immature neutrophils express more CD184 than mature neutrophils, to promote their retention in the bone marrow. But as they age, expression of CD184 increases again favoring return to the bone marrow for clearance <u>59</u>. Here we identified a CD10^-^CD177^+^ immature neutrophil subset that resembles the CD10^-^CD16^low^CD11b^low^ population which has been recently described in the blood of patients with psoriasis, and shown to express CD184 and have low phagocytic ability and ROS production <u>60</u>. On the other hand, in other inflammatory settings mimicking sepsis (LPS injections in mice), aged neutrophils were shown to be particularly effective in migration to sites of inflammation and to exhibit a higher phagocytic activity as compared with the subsequently recruited non-aged neutrophils <u>61</u>. Though the impact of CD184/aging on phagocytic activity is still unclear, our findings identified an increased expression of CD184 in the CD10^-^ CD177^+^ population, in a subset of sepsis patients (**Figure 3D**). Understanding the functional relevance of increased aging of immature neutrophils may provide interesting insights into predicting patient-specific disease outcomes in sepsis.

Results from our study also indicated an increased expression of HLA-DR in the CD10^-^ CD177^+^ sepsis neutrophils (**Figure 3D**). While the expression and role of HLA-DR on neutrophils remains controversial <u>62,63</u>, the notion that under appropriate inflammatory conditions neutrophils can acquire the ability to present antigen and stimulate (or inhibit) T cell responses has started to emerge <u>64</u>. Cytokines such as GM-CSF, IFN-γ, IL-4, and TNF-α have been shown to induce antigen presenting cell (APC) function in both mature and immature neutrophils <u>65,66</u>. Induction of HLA-DR on neutrophils was also observed in patients with chronic inflammatory diseases associated with high levels of cytokines such as patients with rheumatoid arthritis or granulomatosis <u>67,68</u>. Thus, it appears that the ability of neutrophils to present antigen might be driven by a highly inflammatory environment <u>69</u>. In fact, GM-CSF, IFN-γ, IL-4, and TNF-α are upregulated in a subset of the sepsis patients in our cohort, together with increased expression of HLA-DR and CD80 (**Figures 1 and 4**). Therefore, our results support emerging evidence in the literature, and point toward a more important role for immature neutrophils in driving and maintaining antigen-specific T cell responses than had previously been appreciated.

Our data also identified a cluster of mature neutrophils (cluster 17, **Figures 1 and 2**) which resembles a recently characterized population of CD62L^dim^/CD11b^bright^ neutrophils, induced by endotoxin challenge or by severe injury in humans <u>70</u>. This population has been shown to suppress human T cell proliferation in coculture experiments. Our results show that this subset also expressed high levels of CD274 (PD-L1) and CD300f. The PD-1–PD-L1 axis is a well-known immune suppression mechanism that induces lymphocyte apoptosis and monocyte dysfunction, as observed in animal models of sepsis and septic patients <u>71,72</u>. Although blocking this axis using an anti-PD-1 antibody improves survival in septic mice <u>73</u>, the resulting excessive neutrophil engagement with T/B cells and inhibition of humoral immune responses has been shown to be detrimental <u>74,75</u>. In addition, a recent publication suggested that increased PD-L1 expression on human neutrophils delays cellular apoptosis by triggering PI3K–dependent AKT phosphorylation to drive lung injury and increase mortality during clinical and experimental sepsis <u>76</u>. CD300f is another inhibitory receptor expressed in myeloid cells, including mast cells and neutrophils <u>77-80</u>. Recent studies in a model of septic peritonitis, show that absence of CD300f or disruption of ceramide-CD300f interaction stimulates neutrophil recruitment to sites of infection and protects mice from septic death <u>81</u>.

In conjunction with flow cytometry, we also observed differential plasma levels of several key cytokines and soluble factors in sepsis patients. Interestingly, upregulation of classic pro-inflammatory cytokines such as TNFα, IL-1β and IL-6 was heterogenous, upregulation of PTX3, Ang-2, Endocan, Gas6, as well as inflammation marker PCT, was a hallmark of our cohort of septic patients (**Figure S2**). These factors have been previously reported to be increased in septic patients and to drive vascular leak and endothelial dysfunction <u>82,83</u>. PTX3 is an acute-phase protein known to increase rapidly in a variety of inflammatory diseases: it recognizes microbial moieties, activates the classical pathway of complement, and facilitates recognition by macrophages and dendritic cells <u>84,85</u>. In recent years PTX3 was described to have a profound effect on the endothelium and induce vascular dysfunction <u>86-88</u> and levels of PTX3 have been shown to predict and correlate with severity of disease and mortality in sepsis <u>89-91</u>.

However, a relationship between these factors and specific immune cell subsets in the context of sepsis has not been discussed previously. Strikingly, the correlation of neutrophil surface marker expression and plasma levels of soluble factors observed in this study revealed that CD10 (a marker of neutrophil maturity) is inversely correlated to these soluble factors (**Figures 4D**). Thus, we can hypothesize that immature neutrophils might be the drivers of vascular inflammation/leak in sepsis patients. Of note, immature neutrophils have been shown to store and release PTX3 during inflammation <u>92-94</u>. Furthermore, though Ang-1/Ang-2 and Endocan have been associated with inflammation in sepsis <u>95-97</u>, and can influence neutrophil functions such as migration and NETosis <u>98,99</u>, their relationship to neutrophil subsets in sepsis has not been investigated. For the first time, our study demonstrates a significant novel correlation between immature neutrophils and soluble factors that act as drivers of vascular dysfunction.

Sepsis is also characterized by metabolic changes that impact normal metabolic homeostasis. Some of these changes have been suggested as prognostic factors/biomarkers for severity of the disease. For example, lactate, a metabolite of anaerobic metabolism, is used in clinical settings as a marker for sepsis diagnosis. However, the predictive value of this parameter alone remains controversial <u>100-102</u>. In our cohort, (**Table S1**) lactate levels varied from 0.1 to 10, falling outside of the lowest threshold recommended for diagnosis (>2.3 mmol. L^-1^) <u>103,104</u>, and highlighting the challenge of patient heterogeneity. Additional work has examined alterations in metabolite levels associated with sepsis and to date no single compound has shown sufficient sensitivity and specificity to be used as a routine biomarker for early diagnosis and prognosis.

In general, metabolites that belong to the glycolysis, TCA cycle, fatty acid oxidation, and amino acid pathways, play important roles in sepsis and septic shock associated with different causal agents <u>105</u>. Our results (**Figure 5A**) are in line with published work <u>106-109</u> and the metabolic profile used in our study clearly separates healthy subjects from septic patients. One of the novel insights from the current study is a decreased concentration of IPA in septic patients compared to healthy donors. To our knowledge, this trend has not been reported previously. IPA is produced by *Clostridium sporogenes* <u>110</u> in the gastrointestinal tract and has been shown to be a ligand for the pregnane X receptor (PXR) <u>111;</u> consequently, it plays a role in regulating endothelial function <u>112</u>.

Additionally, previous studies have demonstrated IPA’s anti-inflammatory properties <u>113,114</u>, among other therapeutically beneficial effects <u>115,116</u>, which is consistent with the decreased abundance in septic patients observed in this study. Further studies are necessary to better understand the role of IPA in sepsis. In this study, four surface markers out of 14 (CD10, CD16, CD80 and CD86), and 42 metabolites/lipids (out of were significantly different between sepsis and control groups. Interestingly, clustering based around these features revealed that two separate subgroups of septic patients could be clearly discriminated (**Figure 5B**). CD10, whose expression discriminated between mature and immature neutrophils, was the top downregulated surface marker by effect size, inversely correlating with fatty acids and positively correlating with lysoPCs. While several studies report dysfunctional metabolism in sepsis, findings from these studies are observational, not correlative. Our results reveal a novel correlation between expression of CD10 and aberrant levels of plasma metabolites in sepsis patients, thereby indicating an association between immature neutrophils and impaired metabolism. Taken together, integrating data from flow cytometry, cytokine, and metabolite measurements, and performing univariate analysis, our findings reveal a novel link between CD10 expression level, increased vascular permeability, and dysfunctional metabolism in sepsis.

Many of the molecules identified in this study (i.e. PTX3, Ang2, Endocan, PCs, LPA, TNF-α, IL6) have been tested separately as possible diagnostic/prognostic/aiding in patient stratification in sepsis, but none have been successfully adopted. Using multivariate machine learning classifiers, we integrated neutrophil subsets (surface markers), pro/anti-inflammatory and vascular dysfunction mediators and metabolic profiles to identify a multi-omics classifier for sepsis (**Figure 6**). One key finding of our study is that features from each layer were selected by this classifier and that neutrophil surface markers were overrepresented even though that data layer comprised the least number of possible features to select (**Figure 6** and **Figure S3**). This may suggest that a multivariate classifier that utilizes information from several data layers, including the marker panel presented in this paper, may prove valuable for diagnosing sepsis in the future.

Overall, we have developed a systems immunology approach to profile neutrophil signatures and their impact on various aspects of inflammation in sepsis patients. Our approach reinforces the importance of data integration and provides valuable novel insights that will drive future mechanistic studies targeting the trifecta of surface marker expression, cytokines, and metabolites, in sepsis. Specifically, this study identifies an association between CD10^-^ immature neutrophils and markers of endothelial dysfunction indicated by increased PTX3 and dysregulated metabolism suggested by increased FA, in sepsis patients.

## Materials and Methods

### Reagents

Percoll (17-0891-02) was purchased from GE Healthcare Life Sciences. HBSS (14175095), Human IL-2 (PHC0021), RPMI-1640 (A1049101), FBS (10082-139), Penicillin/Streptomycin (15140-122), LIVE/DEAD® Fixable Yellow Dead Cell Stain Kit (L34959), CellTrace CFSE Cell Proliferation Kit (C34554), and Trypan Blue Stain (15250-061) were all purchased from Thermo Fisher. Phytohemagglutinin (PHA) (L1668) and BSA (A9418) were purchased from Sigma.

### Study samples and clinical protocol

Patient cohort comprised subjects with sepsis presenting to the Emergency Department at the Massachusetts General Hospital (MGH) between June 2018 and June 2020. This study was approved by the Institutional Review Boards Harvard and at Partners HealthCare (MA, USA), under protocols 2018P000224. Age- and sex-matched healthy control subjects were collected at MRL. Patients were adjudicated as having sepsis based on Sepsis-3 criteria: Presence of organ dysfunction as a result of bacterial infection. As such, all subjects adjudicated as sepsis received 4 or more days of antibiotic therapy and had an elevated sequential organ failure assessment (SOFA) score upon enrollment. Sepsis severity was identified using the Acute Physiology and Chronic Health Evaluation (APACHE)II score and Sepsis-related Organ Failure Assessment (SOFA) score. Informed consent was obtained from patients or their surrogates. Blood samples from patients and healthy controls were drawn with Na-Heparin Vacutainer tubes (BD Biosciences) and processed within 2 h of collection.

Aliquots of blood from heparin tubes were stained for flow cytometry panels (see below) or centrifuged at 2,000*g* for 10 min and plasma stored at −80 °C. Patient data, including clinical course, relevant laboratory testing, sign of organ disfunction, and 90-day mortality, were collected.

### Neutrophil isolation

Neutrophils were isolated according to a single-step separation procedure with slight modifications <u>117</u>. Blood (5 ml) was layered over a two-step Percoll gradient formed by 4 ml of 75% isotonic Percoll (75% Percoll, 10% PBS 10×, 15 ml of H2O, density (d) 1.103 g/ml, 300–310 mOsM) and 4 ml of 62% isotonic Percoll (62% Percoll, 10% PBS 10×, 28% H2O, d 1.078, 300–310 mOsM) in 15-ml conical test tubes. After centrifugation for 25 min (10 min at 200 × *g* and 15 min at 400 × *g*) at 20°C, neutrophils, located at the interface between the two Percoll solutions, were collected, diluted in Ca^2+^- and Mg^2+^- free HEPES-buffered saline solution containing BSA (HBSS-BSA, 145 mM NaCl, 5 mM KCl, 5 mM HEPES, 5 mM glucose, 0.2% BSA, pH 7.4), and centrifuged for 5 min at 200 × *g*. The procedure was conducted at room temperature and in the absence of divalent cations to prevent neutrophil aggregation and activation. All subsequent experiments were conducted in HBSS.

### Flow cytometry staining and acquisition

Antibodies used for flow cytometry were purchased from BD Biosciences and listed in **Table S2.** Neutrophils were incubated with Fc block (BD Biosciences, 564220) for 10 min prior to staining in staining buffer for 30 min at 4°C. Samples were washed and resuspended in 200ul of staining buffer for acquisition by flow cytometry on a BD FACS Symphony and analyzed with FlowJo (Tree Star) analysis.

### Input data

This study was comprised of three data layers, namely cytokines, neutrophil flow cytometry markers and metabolites. However, all data layers were not able to be collected from all subjects. A table below represents the total number of subjects for each data layer.

### Flow cytometry data analysis pipeline

To analyze and visualize the neutrophil cell surface marker expression via high dimensional flow cytometry (HDFlow), we developed a customized analysis pipeline similar to CyTOF Workflow <u>118</u>. This pipeline consists of a series of steps which are outlined below. The first step in the pipeline was to subset the total number of cells to a smaller, more manageable number (downsampling) and transform the single cell data. We downsampled each sample to 4000 cells. We used the hyperbolic arcsine transformation with a cofactor of 150 to transform the data from an exponential distribution into a normal distribution. The second step in the pipeline was to generate subject-level diagnostic plots, including multidimensional scaling (MDS). We produced this plot using the median transformed expression value for each antigen for each subject as input, and calculated similarity values between pairs of samples which defined the two-dimensional plot showing how similar samples are to each other. The third step in the pipeline was cell clustering. We clustered the cells using a cytometry-specific self-organizing map (SOM) algorithm <u>119</u> called FlowSOM <u>35</u>. The algorithm placed cells into a single node of a 10x10 node grid based on the similarity of that cell to other cells in the node. FlowSOM also performed clustering with ConsensusClusterPlus <u>120</u> on top of the SOM to create a hierarchical clustering of the nodes. Tree cutting was then used to give a specific number of clusters for downstream analysis. We chose 20 initial clusters and then merged six of these 20 clusters into three based on their similar expression patterns, giving us the final 17 clusters. The fourth step in the pipeline was to generate a cell-level Uniform Manifold Approximation and Projection (UMAP) plot <u>36,37</u> and then color cells based upon feature expression. UMAP plots contain all neutrophils in the dataset, colored according to either cluster membership from the FlowSOM step or according to transformed CD10 or CD177 transformed expression. Additional UMAP plots were generated based around the previous coordinates while separating cells from healthy donors and sepsis patients. For differential abundance we utilized a generalized linear mixed model (GLMM) with outcome variables as percentages, while for differential expression we utilized a linear mixed model (LMM) with the outcome variable as median marker expression. In each model, we used patient as a fixed effect and disease status as a random effect. Since the sepsis patients were age- and sex-matched to the healthy controls, we did not include age or sex as covariates in the model. To test coefficients of the model, a log-ratio test was used. To correct for multiple comparisons, we adjusted the nominal p-values from the models using false discovery rate (FDR B&H method).

### Cytokine analysis

Cytokines and chemokines in serum were measured using Magnetic Luminex Screening Assay using the Curiox DropArray System (Curiox Biosystems). The human cytokine/chemokine magnetic bead kit (38-plex) (HCYTOMAG-60K-38X, Millipore) and a custom-made panel (36-plex) (R&D Systems) were used according to manufacturer’s instructions. Data were acquired using xPonent software and represent the median fluorescence intensity of the respective analyte. Sample concentration was calculated by the same software. Each value was measured in duplicates.

### Serum endogenous metabolites and lipids analyses using Biocrates kits

Serum metabolites and lipids were analyzed using a commercial kit Biocrates Quant 500. Briefly, all reagents and consumables including analytical standards, internal standards and sample extraction plates were provided in the kit. Quality control (QC) samples containing expected levels of endogenous analytes were also shipped within the kit. 10μL of each serum sample from the sepsis study was extracted in duplicates along the QCs to generate two extracts separating by polarities of the analytes using different eluents. One extract was analyzed for polar metabolites including amino acids and related, biogenic amines, fatty acids, bile acids, carbohydrates and related, hormones, vitamins, indoles and derivatives, nucleobases and related using liquid chromatography (LC) coupled with triple quadrupole mass spectrometry (QQQ MS). The other apolar extract was analyzed for lipids including acylcarnitines, phosphatidylcholines (PC, including lyso PC), sphingomyelins, cholesteryl esters using flow injection analysis and QQQ MS detection.

A Sciex 6500 QQQ MS equipped with an electrospray ionization source and a Waters Acquity UPLC chromatographic system coupled comprised the LC/MS system. The polar metabolites were analyzed using reversed phase chromatographic separation and scheduled multiple reaction monitoring (sMRM) detection at both positive and negative ionization modes. The lipid analysis was done using single ion monitoring mode on the same QQQ MS. All LC and QQQ MS conditions were provided by Biocrates Quant 500. LC/QQQ MS peak integration was performed using Analyst software (1.6.3).

Quantification and quality control process for both LC/QQQ MS and flow injection/QQQMS analyses were conducted using Biocrates MetIDQ software Qaunt 500 workflow. Lipid species ceramides, diglycerides, and triglycerides in the apolar serum extract were analyzed using a high-resolution mass spectrometer Thermo Scientific Q Exactive HF equipped with HESI-II ion source. Samples were introduced using flow injection following conditions defined within Biocrates p400 kit. The mass resolution was set as 120,000 at m/z400. Data acquisition was performed as full MS scan at positive detection mode. The data process and quality control were completed using Biocrates MetIDQ p400 workflow.

### Normalization and Standardization for Univariate and Multivariate Testing

To satisfy the assumptions of linear regression, each data layer was transformed and standardized as described below. The cytokine data was log2 transformed (log2(x+1)), the metabolites were transformed with variance stabilization transformation (normalize VSN function from LIMMA package <u>121</u> and neutrophil flow cytometry marker expression data was transformed using the following functional form.

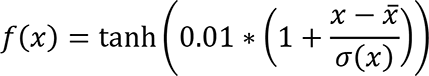

Here *x̅* represents the mean and sigma the standard deviation of a given feature. After transformation, inspection of both histograms and qqplots indicated that transformed data satisfies the assumption of linear regression. Each data set was then standardized using the z-score to have data on the same scale.

### Samples used for each analysis

For analyses performed that combined data layers, we used the intersection of subjects containing relevant data layers. The table below represents the various analyses performed and combinations used.

### Univariate analysis

We defined the effect size for a given feature as the difference in mean transformed and normalized values computed over samples present in the two cohorts. Univariate hypothesis testing using the Student’s *t*-test was performed to obtain the significant p-value associated with the effect size for a given feature. P-values were corrected for multiple hypothesis testing using the False Discovery Rate method (FDR B&H method).

### Multivariate machine learning methods

We built a predictive model for sepsis by examining features in all data layers (flow cytometry, cytokines, and metabolites/lipids) individually (single-omics analysis) to determine which layers had the most predictive signal. The predictive models were constructed in two steps. First, we performed feature selection (FS) using the least absolute shrinkage and selection operator (LASSO) regression <u>122</u> approach followed by Random Forest (RF) <u>123</u> to obtain the model on the selected features.

A stratified nested cross-validation (NCV) strategy <u>124</u> was implemented to minimize overfitting of the predictive model. Data was stratified in such a manner that the proportion of subjects in each cohort was approximately maintained in each fold. We performed 1000 rounds of NCV loops. The outer cross-validation (CV) loop comprised of five folds while the inner loop consisted of three folds. The different numbers of *k*-folds in the inner and outer loops were chosen due to limited number of samples. FS (LASSO regression along with its parameter estimation) and model building steps was performed on the inner loop. Only the features obtained from the FS step were used to build an RF model (using default setting of the ‘random forest’ package in R). The predictive scores of the classifier were obtained on the held out outer CV samples (never seen in the feature selection or the model building steps). We reported the average predictive accuracy and area under the receiver operating curve (AUROC) over 1000 runs of NCV. For multi-omics analysis, all data layers were concatenated. Except for feature selection, we employed all the steps described for the single-omics data on the collated heterogeneous data. Weighted LASSO regression <u>122</u> was used to identify informative features in the multi-omics analysis. These weighting factors were used as a penalization factor in the regularization procedure of weighted LASSO. Lower weights (i.e., less penalty) increased the chances of the feature to be selected in the L1 regularized optimization step. For all features in a data layer, we computed the weights *c*_*ij*_ as the fraction of variation of a given single omics layer, quantified by the mean interquartile range (IQR) of all features divided by the total variation, computed as sum of individual mean IQR of each single omics layer).

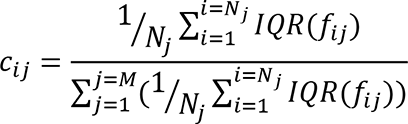

Here *f*_*ij*_ represents the i^th^ feature in the j^th^ dataset, IQR is a function that computes the interquartile range for a given feature, Nj is the total number of features in the j^th^ dataset and M is the total number of datasets in the study. An assumption for this analysis is that large mean IQR computed over of all features in given dataset is undesirable. The weights assigned to cytokine R&D panel, cytokine Millipore panel, flow cytometry panel and the metabolomics panel were 0.20, 0.22, 0.31 and 0.27 respectively. The final panel of features was selected based on the average value of beta, obtained in the FS step, over 1000 runs of NCV. The beta values corresponded to the coefficients obtained when one fits a LASSO model to the data. Average beta values in this analysis ranged from a minimum value of 0 to maximum value of 1.58. For the multi-omics analysis, we reported all features with average beta greater than 0.01.

### Packages used

Heatmaps and correlograms were generated using the function heatmaps2 in the gplots package <u>125</u> in R v3.6.3, while volcano plots were generated using the ggplot2 package <u>126</u> in R v3.6.3 <u>127</u>. The Student’s *t*-test, FDR correction, and Fisher’s exact test were performed using base R, and the Wilcoxon test was done using the R package ggpubr <u>128</u>.

## Supporting information

Correlation Table

## Supplementary Material

**Table S1.**
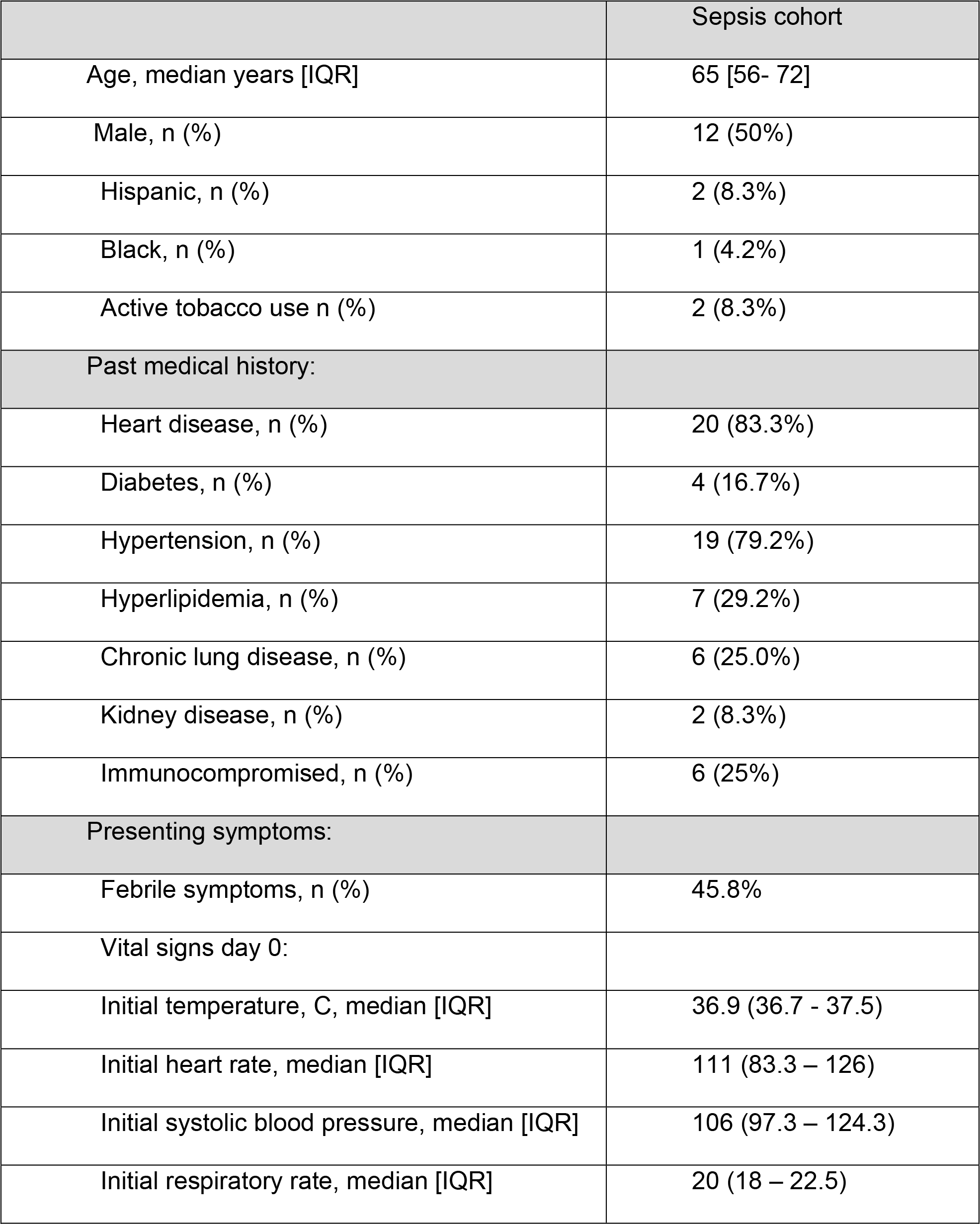
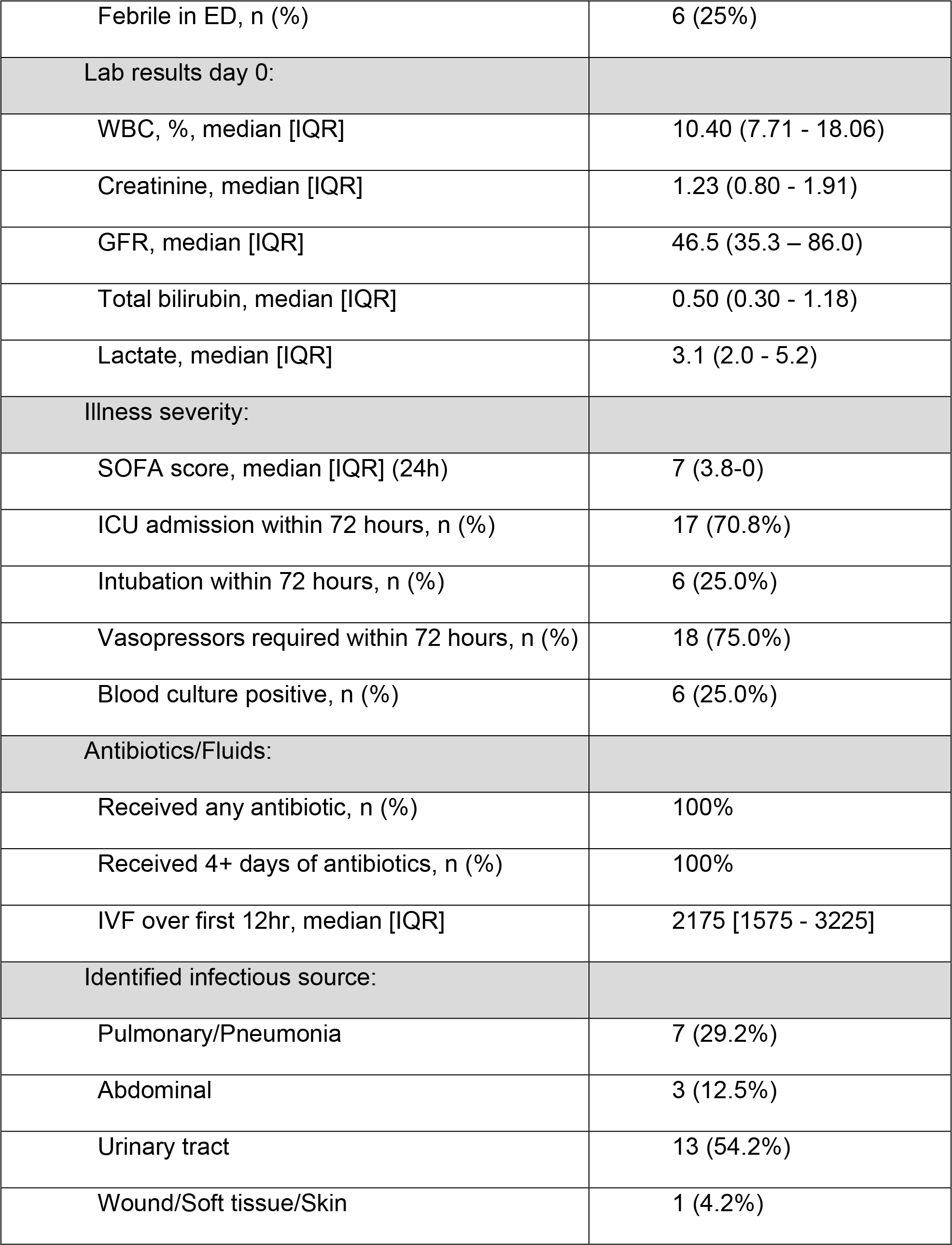
Patient characteristics by infection status.

|  | Sepsis cohort |
| --- | --- |
| Age, median years [IQR] | 65 [56- 72] |
| Male, n (%) | 12 (50%) |
| Hispanic, n (%) | 2 (8.3%) |
| Black, n (%) | 1 (4.2%) |
| Active tobacco use n (%) | 2 (8.3%) |
| Past medical history: |  |
| Heart disease, n (%) | 20 (83.3%) |
| Diabetes, n (%) | 4 (16.7%) |
| Hypertension, n (%) | 19 (79.2%) |
| Hyperlipidemia, n (%) | 7 (29.2%) |
| Chronic lung disease, n (%) | 6 (25.0%) |
| Kidney disease, n (%) | 2 (8.3%) |
| Immunocompromised, n (%) | 6 (25%) |
| Presenting symptoms: |  |
| Febrile symptoms, n (%) | 45.8% |
| Vital signs day 0: |  |
| Initial temperature, C, median [IQR] | 36.9 (36.7 - 37.5) |
| Initial heart rate, median [IQR] | 111 (83.3 – 126) |
| Initial systolic blood pressure, median [IQR] | 106 (97.3 – 124.3) |
| Initial respiratory rate, median [IQR] | 20 (18 – 22.5) |
| Febrile in ED, n (%) | 6 (25%) |
| Lab results day 0: |  |
| WBC, %, median [IQR] | 10.40 (7.71 - 18.06) |
| Creatinine, median [IQR] | 1.23 (0.80 - 1.91) |
| GFR, median [IQR] | 46.5 (35.3 – 86.0) |
| Total bilirubin, median [IQR] | 0.50 (0.30 - 1.18) |
| Lactate, median [IQR] | 3.1 (2.0 - 5.2) |
| Illness severity: |  |
| SOFA score, median [IQR] (24h) | 7 (3.8-0) |
| ICU admission within 72 hours, n (%) | 17 (70.8%) |
| Intubation within 72 hours, n (%) | 6 (25.0%) |
| Vasopressors required within 72 hours, n (%) | 18 (75.0%) |
| Blood culture positive, n (%) | 6 (25.0%) |
| Antibiotics/Fluids: |  |
| Received any antibiotic, n (%) | 100% |
| Received 4+ days of antibiotics, n (%) | 100% |
| IVF over first 12hr, median [IQR] | 2175 [1575 - 3225] |
| Identified infectious source: |  |
| Pulmonary/Pneumonia | 7 (29.2%) |
| Abdominal | 3 (12.5%) |
| Urinary tract | 13 (54.2%) |
| Wound/Soft tissue/Skin | 1 (4.2%) |

**Table S2.**

| **Neutrophil Panel** | **Target** | **Fluorochrome** | **Clone** | **Brand** | **Cat NO** |
| --- | --- | --- | --- | --- | --- |
|  | CD11b (Mac-1) | FITC | ICRF44 | BD Biosciences | 562793 |
|  | CD66b | PerCP-Cy5.5 | G10F5 | BD Biosciences | 562254 |
|  | CD63 | Alexa 647 | H5C6 | BD Biosciences | 561983 |
|  | CD45 | Alexa 700 | HI30 | BD Biosciences | 560566 |
|  | CD62L | APC-Cy7 | DREG-56 | Biolegend | 304814 |
|  | CXCR4 (CD184) | BV 711 | 12G5 (RUO ) | BD Biosciences | 740799 |
|  | CD11c | BV510 | B-Ly6 | BD Biosciences | 563026 |
|  | CD10 | BUV737 | HI10a | BD Biosciences | 564959 |
|  | CD16 | PE | 3G8 | BD Biosciences | 555407 |
|  | CD300f | BUV395 | UP-D1 | BD Biosciences | 745678 |
|  | CD80 | BV786 | L307.4 | BD Biosciences | 564159 |
|  | CD86 | PE-CF594 | 2331 | BD Biosciences | 562390 |
|  | CD177 | BUV421 | Mem166 | BD Biosciences | 564240 |
|  | CD274 | PE Cy7 | MIH1 | BD Biosciences | 558017 |
|  | HLA-DR | BV650 | G46-6 | BD Biosciences | 564231 |

**Table S3.** for samples for different data layers.

| **Data Layer** | **No. of Samples:**<br>**Sepsis** | **No. of Samples:**<br>**Controls** | **Total No. of**<br>**Samples** |
| --- | --- | --- | --- |
| Cytokines | 24 | 19 | 43 |
| Flow Cytometry | 24 | 19 | 43 |
| Metabolites | 14 | 11 | 25 |

**Table S4.** for samples used in analyses performed.

| **Analyses** | **No. of**<br>**Samples:**<br>**Sepsis** | **No. of**<br>**Samples:**<br>**Controls** | **Total No. of**<br>**Samples** | **Analysis**<br>**Method** |
| --- | --- | --- | --- | --- |
| Flow Cytometry | 24 | 19 | 43 | Univariate |
| Cytokines | 24 | 19 | 43 | Univariate |
| Metabolites | 14 | 11 | 25 | Univariate |
| Flow Cytometry<br>and Cytokines | 21 | 17 | 38 | Univariate |
| Flow Cytometry<br>and Metabolites | 12 | 10 | 22 | Univariate |
| Cytokines, Flow<br>Cytometry, and<br>Metabolites | 12 | 9 | 21 | Multivariate<br>Machine<br>Learning |

**Supplemental Figure S1.**
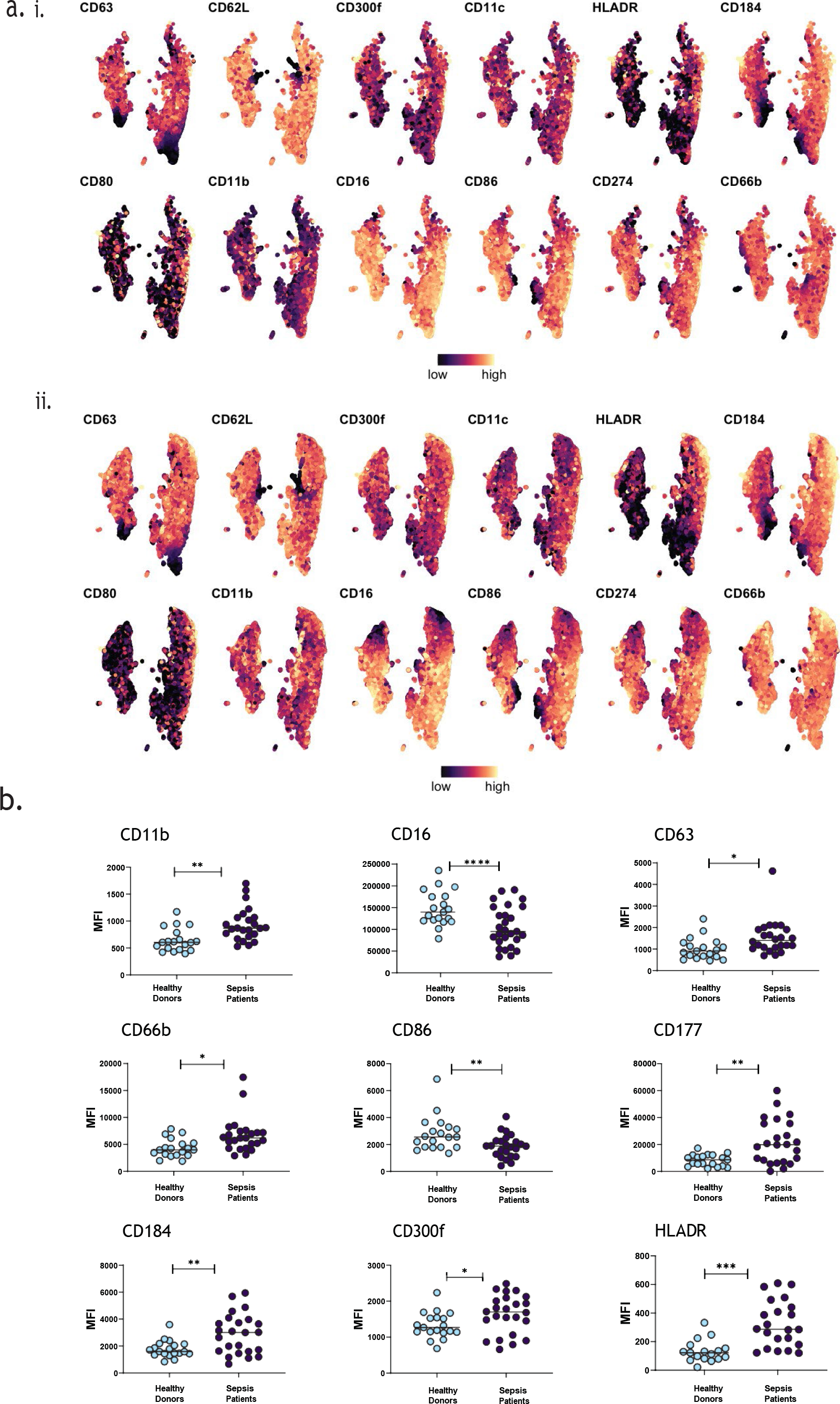
Individual neutrophil marker expression in cell clusters. **(A)**Neutrophil surface marker UMAP from Figure 1C split by marker and by cohort (sepsis patients or healthy donors). Points in the UMAP were colored according to marker expression, from low (dark blue) to high (bright yellow). **(B)** Differential expression of surface markers in sepsis patients (n=24) vs healthy controls (n=19) measured by flow cytometry.

**Supplemental Figure S2.**
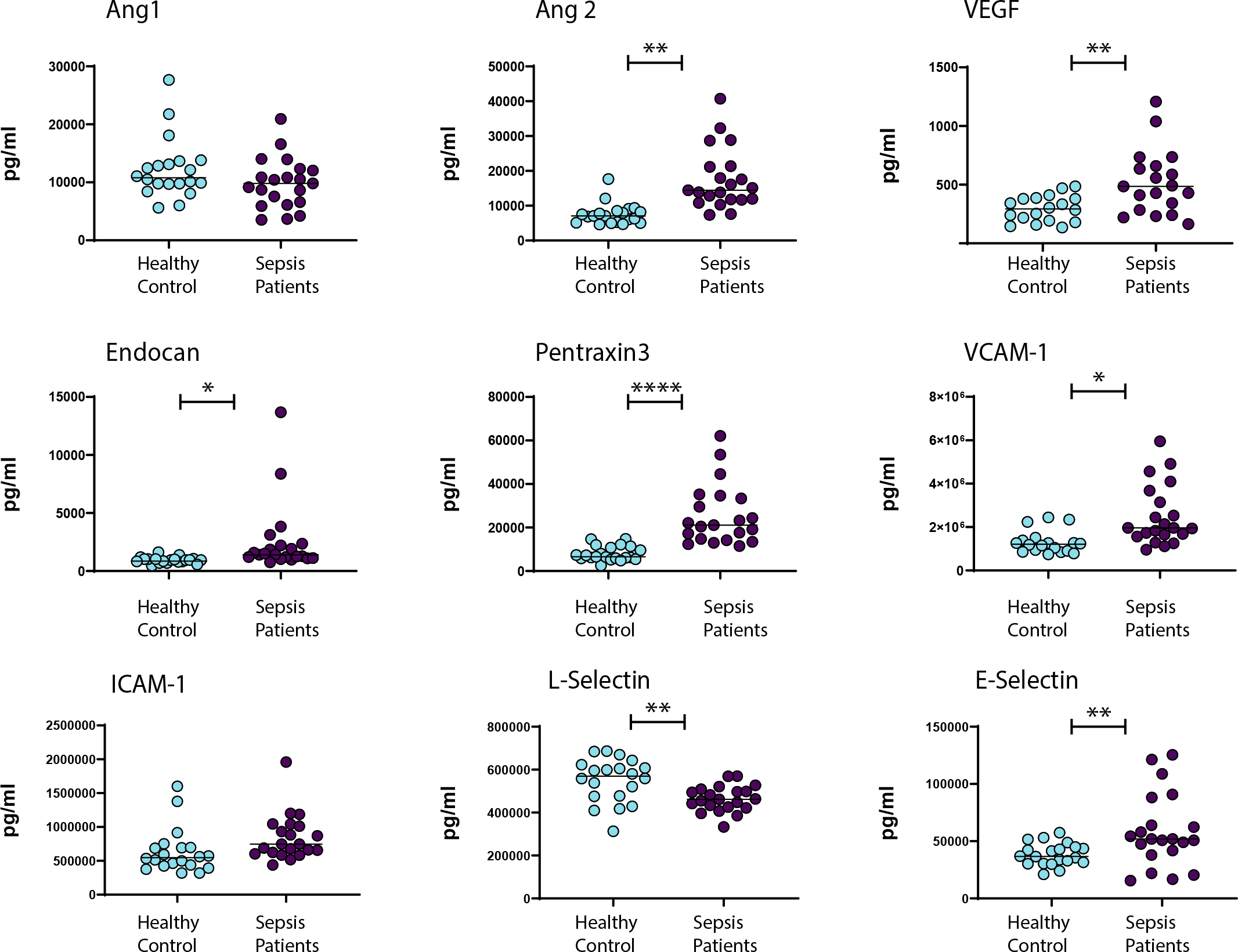
Differential Expression of cytokines and soluble factors in sepsis patients. Differential expression of soluble factors indicative of vascular dysfunction in sepsis patients (n=24) vs healthy controls (n=19) measured by Magnetic Luminex Screening Assay using the Curiox DropArray System.

**Supplemental Figure S3.**
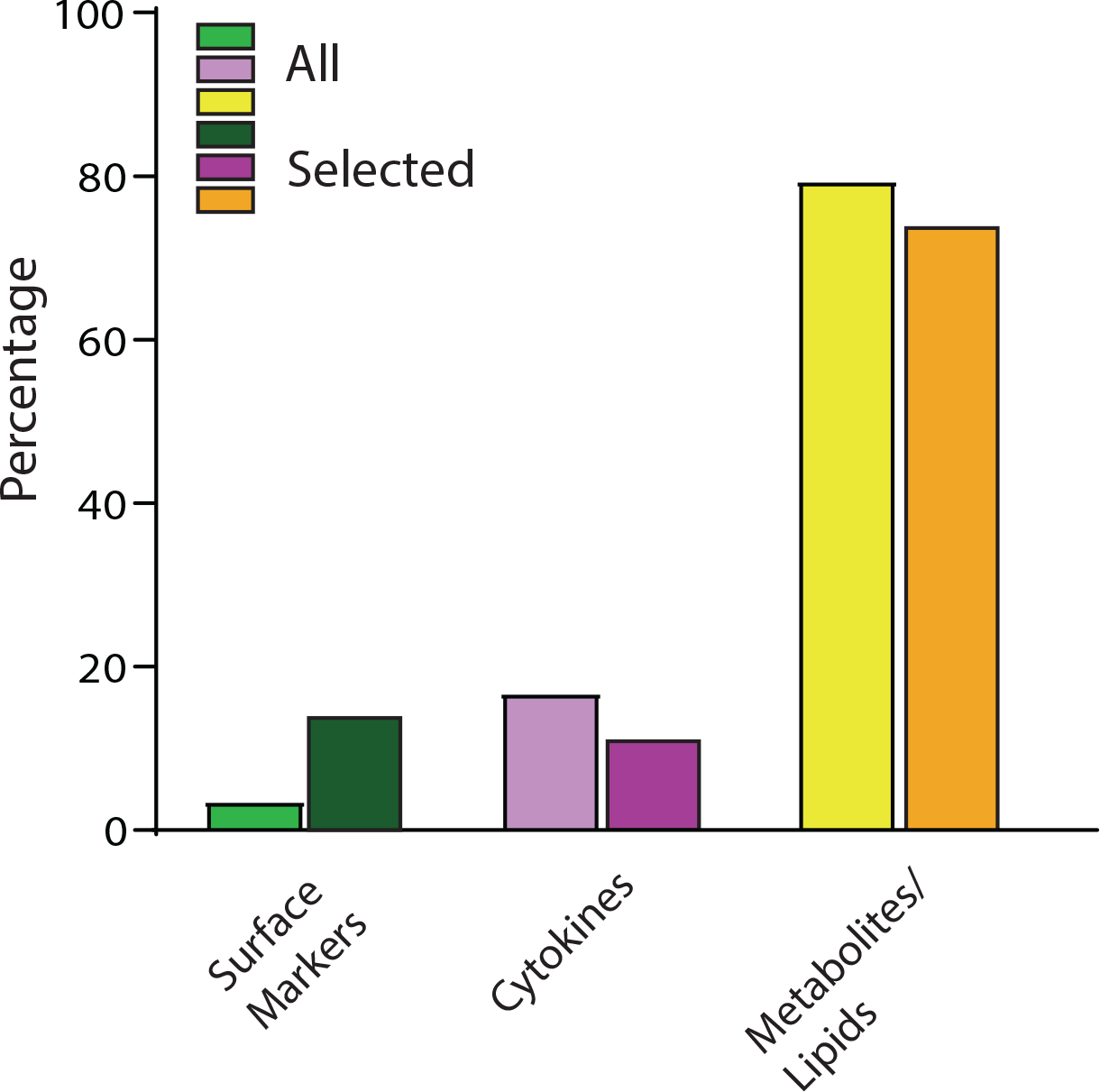
Feature percentages selected for classifier. Bar graph showing the percentage of features from all data layers (left bar), and the corresponding percentage of features selected by the classifier (right bar).

## Notes

### Competing Interest Statement

The authors have declared no competing interest.

